# The role of the gustatory cortex in incidental experience-evoked enhancement of later taste learning

**DOI:** 10.1101/369520

**Authors:** Veronica L. Flores, Tamar Parmet, Narendra Mukherjee, Sacha Nelson, Donald B. Katz, David Levitan

## Abstract

The strength of learned associations between pairs of stimuli is affected by multiple factors, the most extensively studied of which is prior experience with the stimuli themselves. In contrast, little data is available regarding how experience with *incidental* stimuli (independent of any conditioning situation) impacts later learning. This lack of research is striking given the importance of incidental experience to survival. We have recently begun to fill this void using conditioned taste aversion (CTA), wherein an animal learns to avoid a taste that has been associated with malaise. We previously demonstrated that incidental exposure to salty and sour tastes (taste pre-exposure—TPE) enhances aversions learned later to sucrose. Here, we investigate the neurobiology underlying this phenomenon. First, we use immediate early gene (c-Fos) expression to identify gustatory cortex (GC) as a site at which TPE specifically increases the neural activation caused by taste-malaise pairing (i.e., TPE did not change c-Fos induced by either stimulus in isolation). Next, we use site-specific infection with the optical silencer Archaerhodopsin-T to show that GC inactivation during TPE inhibits the expected enhancements of both learning and CTA-related c-Fos expression, a full day later. Thus, we conclude that GC is almost certainly a vital part of the circuit that integrates incidental experience into later associative learning.

## INTRODUCTION

Consistent pairing of specific taste stimuli with strong reinforcement leads animals to adapt their future responses to those stimuli, thereby making them more successful at consuming nutrients and avoiding toxins. In the lab, the most well-known variety of this adaptive process is called conditioned taste aversion (CTA), wherein animals learn to avoid a taste conditioned stimulus (CS) that has been paired with malaise-inducing unconditioned stimulus (US). While complex, CTA is known to involve: 1) changes in a brainstem-amygdalar-cortical circuit (Bures et al. 1998; Grossman et al. 2008); and 2) synaptic plasticity governed by the degree of the association between the CS and US (Garcia et al. 1966; Revusky 1968; Nachman 1970; Ahlers and Best 1971; Kalat 1971; Balsam et al. 2002; Frankland et al. 2004; Molet and Miller 2014; Adaikkan and Rosenblum 2015; Kirkpatrick and Balsam 2016).

Of course, reliable pairings of stimulus and reward are quite rare in the ongoing stream of experience. Most taste stimuli are seldom experienced with strong reinforcement--they are "innocuous", meaning they occur incidentally. Nonetheless, these experiences are important for survival, in that ostensibly innocuous stimuli can have a measurable impact on behavioral adaptations caused by learning—that is, on associative memory strength (Walters and Byrne 1983; Fanselow and Poulos 2005; Johansen et al. 2011; Kandel et al. 2014). An extensive body of research has shown, for instance, that CTA memory strength is decreased by unreinforced pre-exposure to the CS or US, which renders the stimuli familiar and less salient (e.g., Lubow and Moore 1959; Lubow 1973; Cannon et al. 1975; Lovibond et al. 1984), and pinpoints possible neural loci of these effects (Weiner 2010).

This work leaves unstudied, however, the potential impact of the most common stimuli—those other than the CS and US in some eventually-experienced learning paradigm. There are at least two reasons why it is reasonable to ask whether even totally “incidental” experience with a set of tastes might in fact have an impact on learning about a new taste: 1) general environmental “enrichment” has been shown to affect both neural development and specific sensory responses (Alwis and Rajan 2013; Liu and Urban 2017); and 2) “innocuous” stimuli have been suggested to enhance sub-threshold learning experiences (Ballarini et al. 2009) and latent inhibition (Merhav and Rosenblum 2008). Still, virtually no work has explicitly investigated how incidental taste experience might change the function of CTA learning circuits in the brain. This noticeable gap in the literature is a potentially significant limiting factor on our ability to generalize the results of lab experiments to the human condition—incidental taste experience is omnipresent in the natural world, a fact that stands in stark contrast to the laboratory, in which learning experiments are performed on animals that have never tasted anything but (mild, nearly tasteless) chow.

We have recently begun an inquiry into this topic (Flores et al. 2016), showing that experience with salty and sour tastes (hereafter “taste pre-exposure” or TPE) enhances aversions toward a novel taste; experience with both tastes is more effective than experience with either alone, and three sessions of TPE is more effective than two. These results, which contrast with both classic interference effects that reduce conditioning strength (Pavlov 1927; Bouton 1993; Kwok et al. 2012) and the above-mentioned effects that occur across an entirely different time scale (Riege 1971; Donato et al. 2013; Leger et al. 2015), demonstrate that benign experience with one set of tastes over two to three days can impact the strength of learning established in a later associative conditioning paradigm using a different taste.

While much remains to be learned about this behavioral phenomenon, the above results do enable us to formulate basic hypotheses regarding how sensory taste information acquired during TPE is processed in the brain and integrated into future learning. An appropriate place to start in this regard is with primary gustatory cortex (GC), which resides in the anterior insula: GC has been amply shown, in electrophysiological studies (Katz et al. 2001; Grossman et al. 2008; Moran and Katz 2014; Sadacca et al. 2016), immediate early gene imaging (Desmedt et al. 2003; Contreras et al. 2007; Inberg et al. 2013; Uematsu et al. 2015), and loss of function experiments (Berman and Dudai 2001; Stehberg and Simon 2011; Levitan et al. 2016a; Levitan et al. 2016b; Li et al. 2016) to play a role in mediating taste behavior and CTA learning, as well as taste novelty/familiarity (Gallo et al. 1992; Rosenblum et al. 1993; Koh et al. 2003; Bahar et al. 2004; Koh and Bernstein 2005; Roman and Reilly 2007; Merhav and Rosenblum 2008; Nunez-Jaramillo et al. 2008; Lin et al. 2012c; Inberg et al. 2013; Hadamitzky et al. 2015). Here, we test the hypothesis that GC is vital for representing incidental taste experience and for driving the impact of that experience on later CTA learning.

Specifically, the current study: 1) directly characterizes CTA-training-related c-Fos expression—an immunohistological marker for neural activation—in GC following TPE; 2) replicates the previously-shown TPE phenomenon in virus-infected rats; and 3) tests the impact of optogenetic silencing of GC activity during TPE on later learning. Our results support the hypothesis that GC plays an important role in integrating incidental taste experience with future learning (without changing processing of the taste experience alone), thereby mediating later memory formation—both TPE itself, and manipulation of GC specifically during TPE, impact a future CTA (and learning-related neural activation) towards a novel taste.

## RESULTS

### Experiment 1: TPE increases CTA-related c-Fos activity in GC

Rats were subjected to the TPE protocol—3 days of pre-exposure to sodium chloride (N, NaCl, salty taste), citric acid (C, sour taste), and distilled water (W) via IOC—and then given 1 day of aversion conditioning to a novel sucrose CS (Figure 1); TPE in this protocol was previously shown to enhance the strength of CTA (Flores et al. 2016; see Experiment 2 for a replication and extension of the behavioral phenomenon).

**Figure 1.**
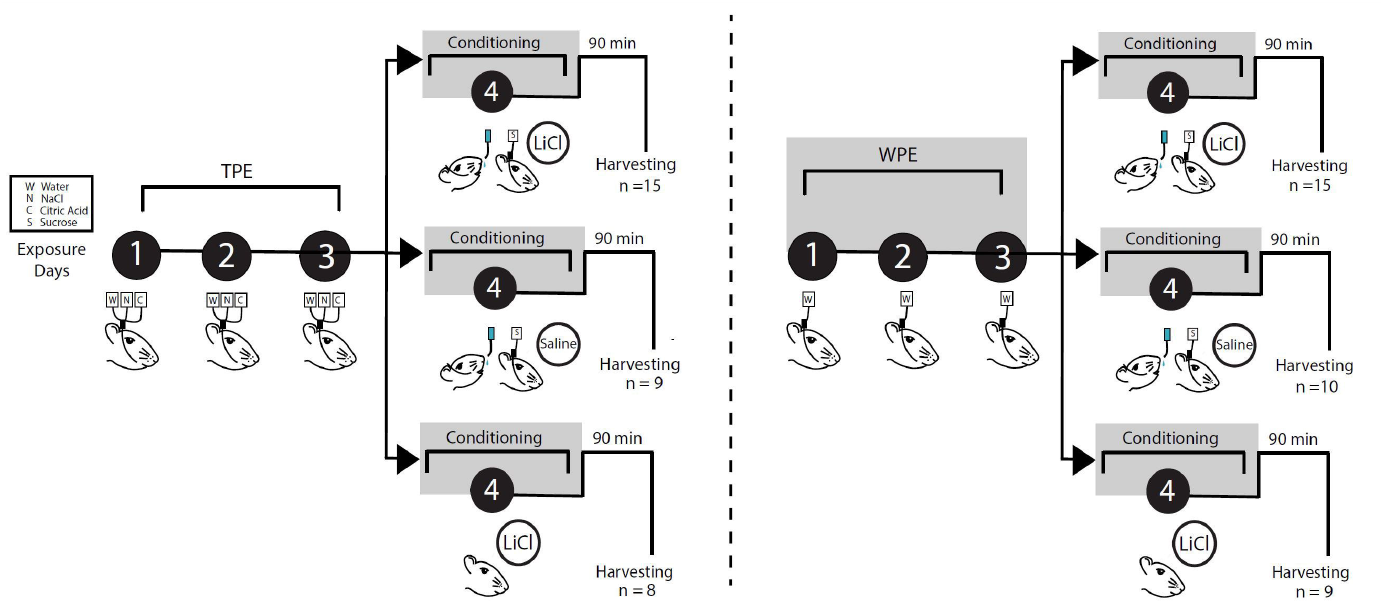
Taste Pre-Exposure paradigm. A complete timeline of the Taste pre-exposure paradigm showing all groups. Animals were divided into 2 groups: TPE (left) or WPE (right), then further divided into three conditioning conditions: sucrose + LiCl, sucrose + saline, and LiCl alone. Schematic demonstrates the four-day experimental paradigm in which rats receive TPE to water (W), sodium chloride (N) and citric acid (C), *via* IOC infusions to the tongue for 3 d (black circles days 1–3). WPE rats underwent 3 identical days of exposure to water. Aversions were then conditioned on the fourth day, when exposure to sucrose (S) is immediately followed by LiCl injections (0.3 M, 0.5% of current body weight), equal dosages of saline or equal dosages of LiCL without sucrose exposure. Control rats were either given saline injections or LiCl alone. Ninety minutes after the conditioning session, rats were perfused for harvesting of gustatory cortex.

Brains were harvested for immunohistochemistry 90 min after the training trial and processed for c-Fos. The 90-min waiting period allowed c-Fos activity to approach its peak (Chaudhuri et al. 2000) consistent with previous studies (Koh and Bernstein 2005; Uematsu et al. 2015), and the decision to look at c-Fos caused by the training trial (the only session that was identical for all trained rats) allowed us to specifically compare the number of GC neurons activated by the associative learning situation in TPE and non-TPE rats. C-Fos differences observed after testing sessions would have been difficult to interpret, as they would reflect some combination of: 1) learning differences; 2) retrieval differences; and 3) differences in consumption caused by differential learning (see Methods).

Given that CTA learning has repeatedly been associated with increased levels of c-Fos expression in GC (Koh and Bernstein 2005; Hadamitzky et al. 2015; Soto et al. 2017), and that TPE enhances learning (Flores et al. 2016), we hypothesized that TPE would enhance learning-related c-Fos expression in GC. We therefore compared c-Fos in CTA-conditioned TPE rats to that of five controls: 1) rats that underwent the exact same CTA – conditioning procedure, but were pre-exposed only to water (i.e., WPE rats); 2-3) rats that received TPE or WPE, but for which the “training trial” was actually a “sham-training” trial in which sucrose was delivered was paired with harmless saline injection; and 4-5) rats that received TPE or WPE followed by “training” that consisted of LiCl alone (Figure 1). These experiments allowed to test, most centrally, whether CTA training causes more c-Fos activation when preceded by TPE than when preceded by lack of TPE, and whether CTA training causes more c-Fos activation than either sham-training or LiCl alone when each was preceded by TPE. In addition, examining “conditioning-trial” c-Fos in this set of conditions allowed us to test whether sham-training or LiCl alone causes more c-Fos activation when preceded by TPE than when not.

Figure 2B shows, at two magnifications, representative examples of c-Fos expression in GC (Figure 2A, from left to right) for TPE-CTA, WPE-CTA, TPE-Sham and TPE-LiCl (the groups involved in the primary pair of tests above). In the main panels of Figure 2B, black spots represent the somae of c-Fos positive neurons revealed by our immunostaining and imaging procedures (the insets confirm that the signal comes from cell bodies rather than noise). It can easily be seen that TPE led to higher levels of CTA-related (i.e., following Sucrose + LiCl) activation than did WPE, and that even fewer neurons were activated by the taste of sucrose or LiCl alone, despite their being preceded by TPE.

**Figure 2.**
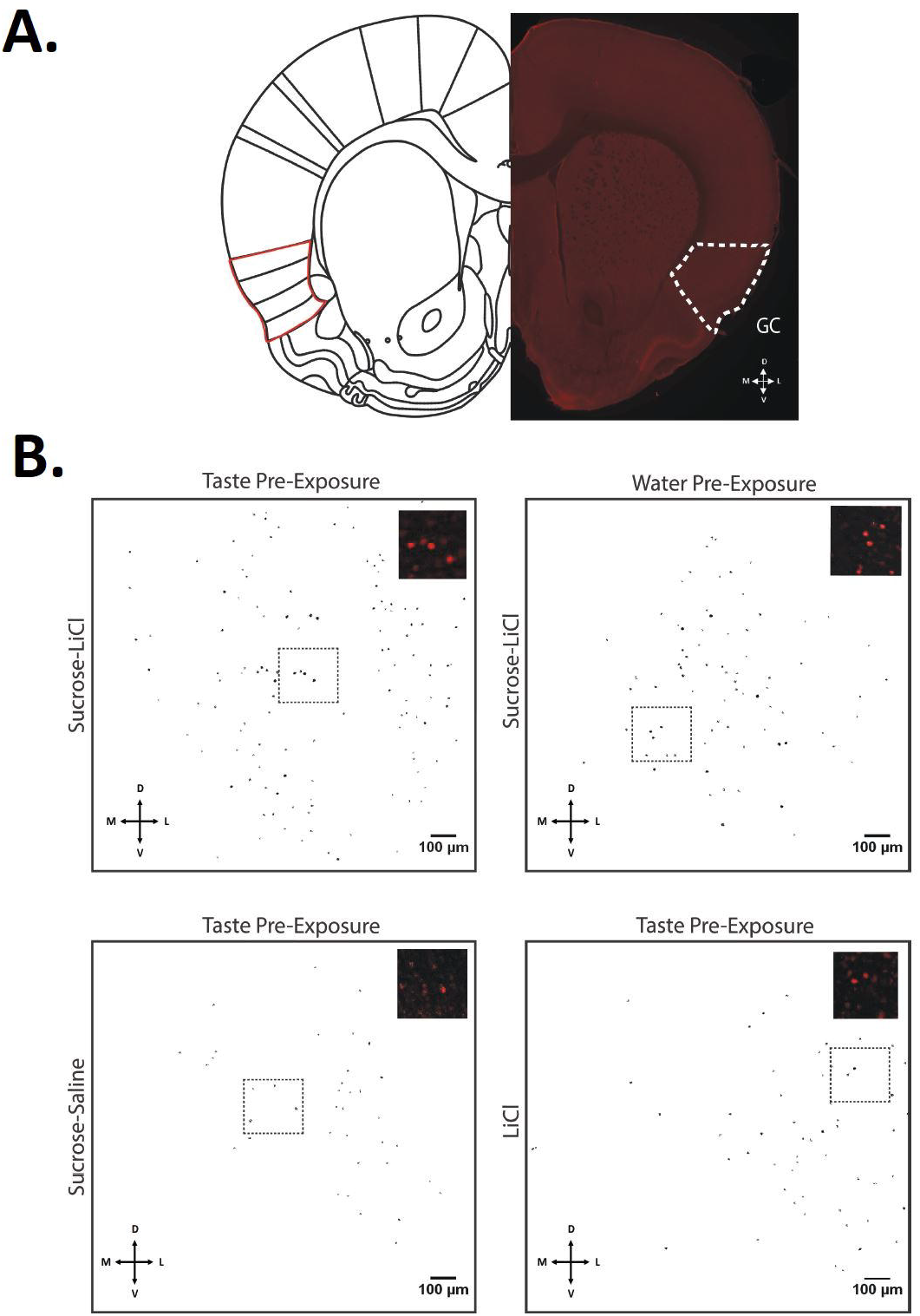
c-Fos positive cells in GC after CTA conditioning to novel sucrose. **A.** A representative coronal slice indicating the location of gustatory cortex (GC, Left-half hemisphere schematic; Paxinos and Watson 2007; permission requested). Bottom directional indicates direction of tissue location—dorsal (D), ventral (V), medial (M), and Lateral (L). **B.** Representative images of c-Fos positive somae (masked in black) in GC for the 4 groups most relevant to the central Experiment 1 hypothesis, quantified by the FIJI Analyzing Particles tool. From top left to bottom right: TPE followed by pairing of sucrose and LiCl; WPE followed by pairing of sucrose and LiCl; TPE followed by a pairing of sucrose and saline; and TPE followed by LiCl alone. Insets represent higher-magnification samples of c-Fos positive somae (red) sampled from the region in the dotted black rectangle (note: quantification took place across the entire masked image).

Analysis of the group data, which are shown in Figure 3A, supports the conclusions suggested by visual scrutiny of the representative data shown in Figure 2B. An ANOVA (see methods) revealed significant differences between the groups (*F* (5, 65) = 5.092, *p* = 0.001); subsequent Fisher’s LSD post hoc analyses reveal a specific increase in CTA-related activation of GC caused by TPE—c-Fos expression for this group (102 ± 8 somae) was higher than the CTA group preceded by WPE (78 ± 9 somae, *p* = 0.025). In fact, CTA preceded by TPE was found to induce significantly higher c-Fos counts than any of the control groups (*p’s all* < 0.05). This supports our central hypothesis that TPE enhanced the processing of sucrose-LiCl pairing.

Of course, as noted in the Introduction, CTA would also be strengthened if TPE was to enhance responsiveness to either the taste or LiCl itself (Logue 1979; Franchina and Slank 1988; Flores et al. 2016), perhaps by enhancing novelty or neophobia. The c-Fos evidence suggests that TPE does not enhance processing of either stimulus presented alone, however: levels of c-Fos expression for TPE-sucrose (+ saline) and TPE-LiCl groups proved indistinguishable from those observed for WPE-sucrose (Fisher’s LSD, *p* = 0.596) and WPE-LiCl (Fisher’s LSD, *p* = 0.234) groups (Figure 3); if TPE had enhanced stimulus novelty, it would have increased c-Fos expression following exposure to sucrose or LiCl (Koh et al. 2003; Lin et al. 2012a).

**Figure 3.**
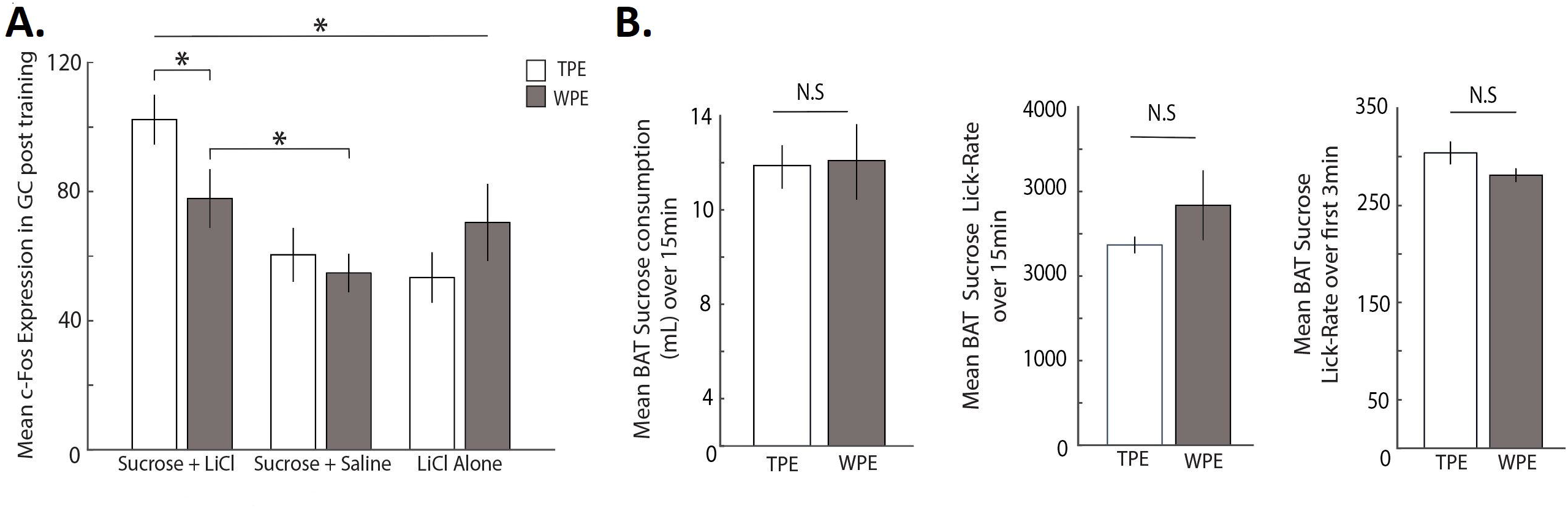
TPE increases CTA-related c-Fos expression in GC. **A.** Pre-exposure to salty and sour tastes (open bars) followed by CTA conditioning resulted in significantly higher c-Fos expression in GC, compared to WPE rats (gray bars); c-Fos in the TPE - CTA conditioning group was also significantly higher than all other groups. These results, and the fact that TPE did not enhance c-Fos in sham and LiCl-alone groups, demonstrate that TPE specifically impacts the pairing of the sucrose with LiCl, and not the processing of either independently. **B. *Left:*** BAT sucrose consumption (mL) across 15-min access to 15mLs sucrose was similar for TPE and WPE rats. ***Middle:*** Lick rate for sucrose was similar for TPE and WPE rats during the entire 15min BAT session. ***Right:*** Initial lick-rate (average licks for the first 3 minutes of the BAT) was similar for TPE and WPE rats. Error bars represent SEM. (*) *p* < 0.05.

As an independent test of this last finding, we collected and performed an analysis of licking to novel sucrose in a brief access task (BAT, Med Associates Inc.). The BAT provides a set of particularly rich, sensitive assays of taste responsiveness using individual licks as the basic units of measure (see Methods and Davis 1989); notably, differential lick rates across even small sets of trials reliably reveal differences in novelty (i.e., neophobia; Lin et al. 2012b; Monk et al. 2014). Our comparison of TPE and WPE rats revealed no such differences in 15-min raw sucrose consumption (*t* (6) = - 0.112, *p* = 0.914) and lick rate averages (*t* (6) = −1.098, *p* = 0.314), or even in the lick rate averages during the initial three minutes of sucrose trials (independent samples t-test, *t* (6) = 1.671, *p* = 0.146, Figure 3B)—in fact, by this most sensitive measure, TPE trends (insignificantly) toward the less novel (a result consistent with previous suggestions about the impact of innocuous taste experience, Capretta et al. 1975; Braveman 1978; Miller and Holzman 1981; Franchina and Gilley 1986; Pliner et al. 1993; Morón and Gallo 2007). Thus, our c-Fos and behavioral results suggest that TPE changes learning itself, rather than enhancing sucrose novelty.

Further examination of the Figure 3A c-Fos data largely conformed to expectation. A significant difference between training-session c-Fos observed in WPE-CTA and WPE-sham rats (Fisher’s LSD, *p* = 0.044) replicated previous studies (Navarro et al. 2000b; Koh and Bernstein 2005; Wilkins and Bernstein 2006; St Andre et al. 2007; Hadamitzky et al. 2015). The difference between the WPE-CTA and WPE-LiCl groups (Fisher’s LSD, *p* = 0.549) failed to achieve significance here, but by far the simplest explanation for this lack of significance, which is of little import in the current study, is the fact that the current protocol was specifically designed to minimize it: the need to avoid a ceiling effect in learning (which would have obscured the enhancement of learning caused by TPE) required that we reduce by half the concentration of LiCl used (see Methods), which in turn ensured that WPE-CTA caused relatively mild conditioning (see Experiment 2 and Nachman and Ashe 1973; Navarro et al. 2000a; Hadamitzky et al. 2015; Flores et al. 2016) that was necessarily difficult to differentiate from the c-Fos induced by a powerful emetic stimulus. Regardless, this result only serves to emphasize the fact that TPE “primes” cortical activation in response to a CTA training trial. This enhancement of neural activation could reasonably be expected to enhance CTA strength.

Finally, we asked one further question with these c-Fos data, examining whether the proposed neural substrate of TPE’s impact on learning might be localized to a specific sub-region of GC. A recent study has demonstrated that lesions large enough to include posterior GC have a larger impact on CTA learning than those limited to anterior GC (Schier et al. 2016); to test whether this anatomical subdivision also offered a more precise characterization of our TPE effect, we re-analyzed c-Fos data from TPE and WPE rats, making three separate measurements for each rat—one in an anterior GC slice, one in a middle GC slice, and one in a posterior GC slice (Figure 4A; see Methods).

**Figure 4.**
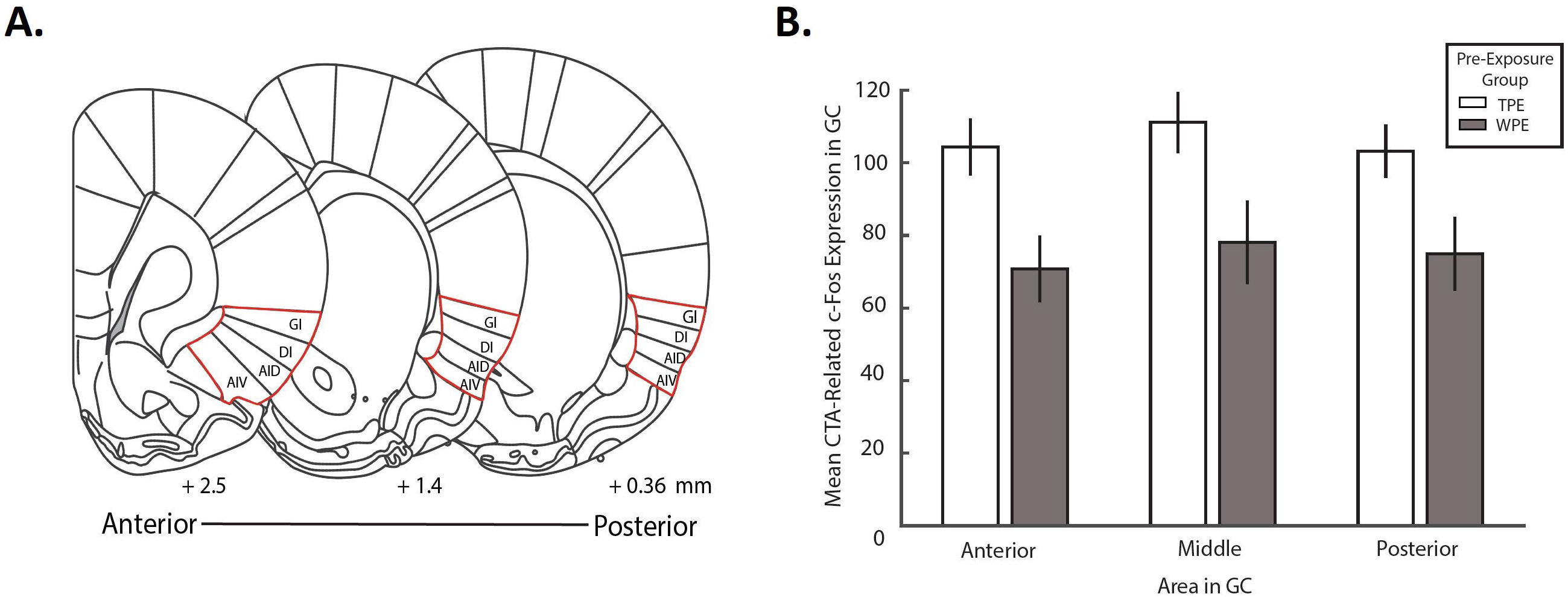
TPE-evoked increases in CTA-related c-Fos expression impacts the entire GC. **A.** Schematics of coronal sections of anterior, middle and posterior regions of GC (Paxinos and Watson 2007; permission requested). Numbers at the bottom indicate distance from bregma (in mm) for designated anterior, middle and posterior regions of GC. Areas outlined in red indicate the locations of gustatory cortex. **B.** Mean CTA-related c-Fos positive somae in GC corresponding to the regions in Panel A for rats given TPE or WPE. The impact of TPE on CTA-related c-Fos expression was similar for anterior, middle or posterior sub-regions of GC. Error bars represent SEM.

The lack of significance in the interaction term of a repeated measures ANOVA (for pre-exposure condition and sub-region) allows us to reject this ancillary hypothesis, revealing that the effect of TPE on CTA-related c-Fos expression did not differ across sub-regions of GC (*F* (2, 52) = 0.168, *p* = 0.846). Nor did the main effect for amount of c-Fos in different GC regions reach statistical significance (*F* (2, 52) = 2.380, *p* = 0.103); only the expected main effect showing that TPE enhances c-Fos was borne out (*F* (1, 26) = 6.783, *p* = 0.015). TPE influenced CTA-related activation across the entirety of GC similarly (Figure 4B).

### Experiment 2: Perturbation of GC activity mitigates the impact of TPE on CTA

The above experiments demonstrate that TPE enhances GC neural responsiveness to the later association of a novel taste with illness. These results suggest, but do not prove, that GC is a part of the circuit responsible for the enhancement of learning caused by TPE (Flores et al. 2016); similarly, they suggest but stop short of conclusively demonstrating a link between the observed enhancement of c-Fos and enhanced learning (although this link has been proposed previously, see Koh and Bernstein 2005; Hadamitzky et al. 2015).

Both of these above hypotheses can be tested with optogenetic silencing of GC. Note, however, that these tests are quite risky and novel. Specifically, if it is true that GC activity during TPE is vital for the subsequent enhancement of learning and learning-related c-Fos, then inhibiting GC activity during TPE should eliminate these enhancements—enhancements observed a full 24 hours after the last session of inhibition. We are unaware of another experiment in which neural inhibition was predicted to have an impact on either learning or neural activation produced by a procedure administered that much later. Here, we test for each of these possible impacts in turn.

#### Experiment 2A (impact of TPE and GCx on learning-related consumption)

To perform a direct test of our hypothesis that GC neural activity plays an important role in the TPE-induced enhancement of learning, we performed a set of experiments in which we optogenetically perturbed GC neural activity (GCx) during TPE sessions using the light-gated optical silencer ArchT (Archaerhodopsin-T, a light activated H+ pump, see Han et al. 2011; Yizhar et al. 2011). The construct was delivered in an adeno-associated virus (AAV) also expressing green fluorescent protein fused to ArchT (AAV9-CAG-ArchT-GFP), allowing us to visualize infection sites (Figure 5A).

**Figure 5.**
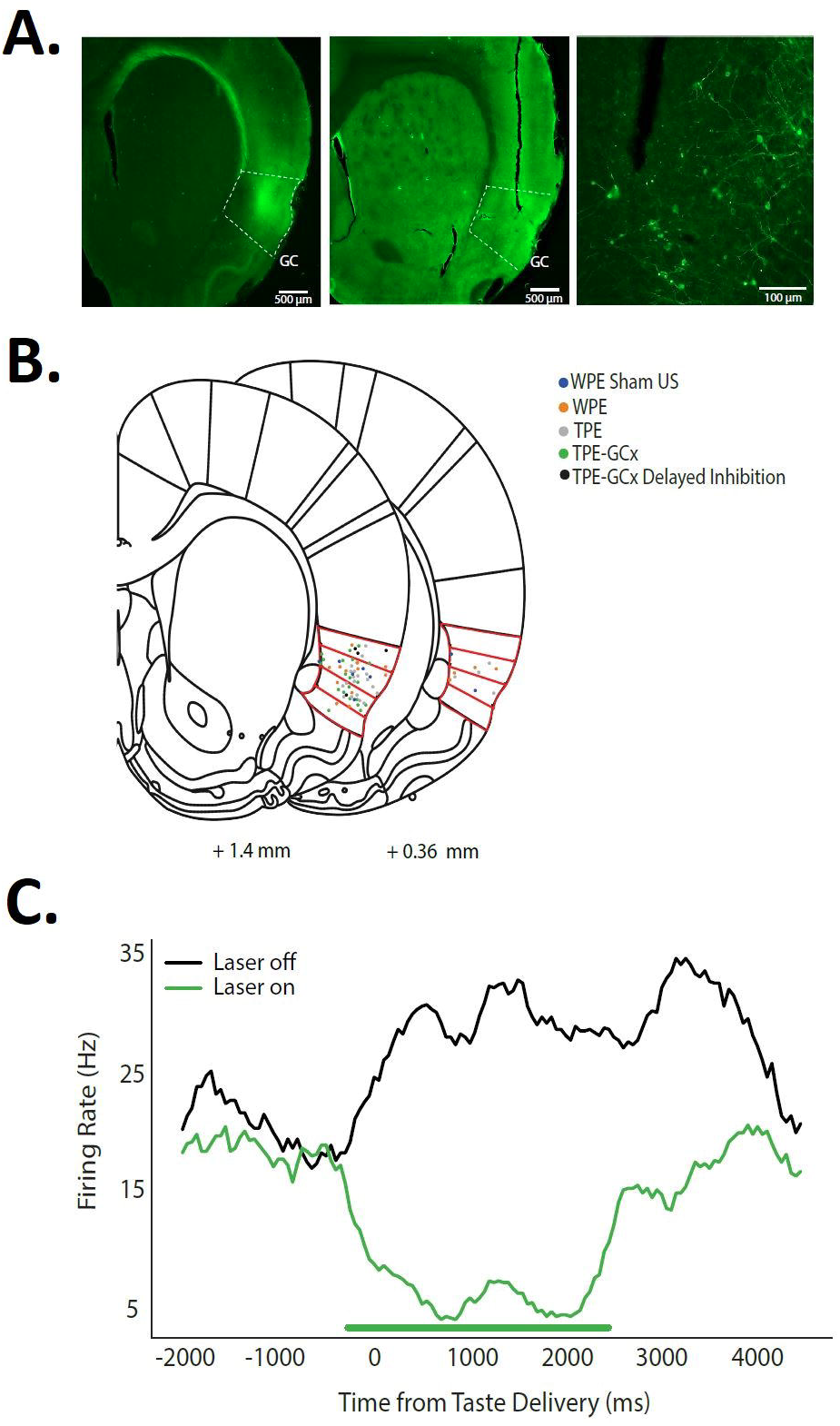
Localization of viral infection and optical fiber placement in GC. **A**. Representative fluorescent images of gustatory cortex confirming infection of ArchT (left) and placement of optical fiber (middle) in GC. The far-right panel shows, at higher magnification, infected neurons in GC stained with GFP at the tip of the fiber track. **B**. Localization of all fiber tips for all optogenetic groups in Experiment 2A, overlain on schematic coronal slices (Paxinos and Watson 2007; permission requested), demonstrating reliable placement in GC; red outlines are the 4 granular regions in GC. Note that for simplicity and demonstration here, fiber localizations for both hemispheres are overlaid on to one. Each rat received 2 fiber depth scores (one per hemisphere), which were then averaged for a single Fiber Depth index per rat (see Methods). **C.** Peristimulus time histogram (PSTH) of a single GC neuron infected with ArchT demonstrating that our optogenetic inactivation protocol has the desired effect. The firing rate of the infected neuron drops drastically while the laser is on (green line, 0-2500ms post taste delivery).

Every rat run in Experiment 2A & B (n = 53; regardless of group) was infected with virus and given fiber optic implants into GC (Figure 5B). Recovery was followed (again, for all rats) by 3 pre-exposure sessions in which GC was illuminated by green (532nm) laser light from 0.85 sec before until 2.5 sec after each fluid delivery, which in GCx rats successfully inhibited GC neuron firing (Figure 5C). The final TPE/WPE day was in turn followed by CTA training and testing sessions (Figure 6).

**Figure 6.**
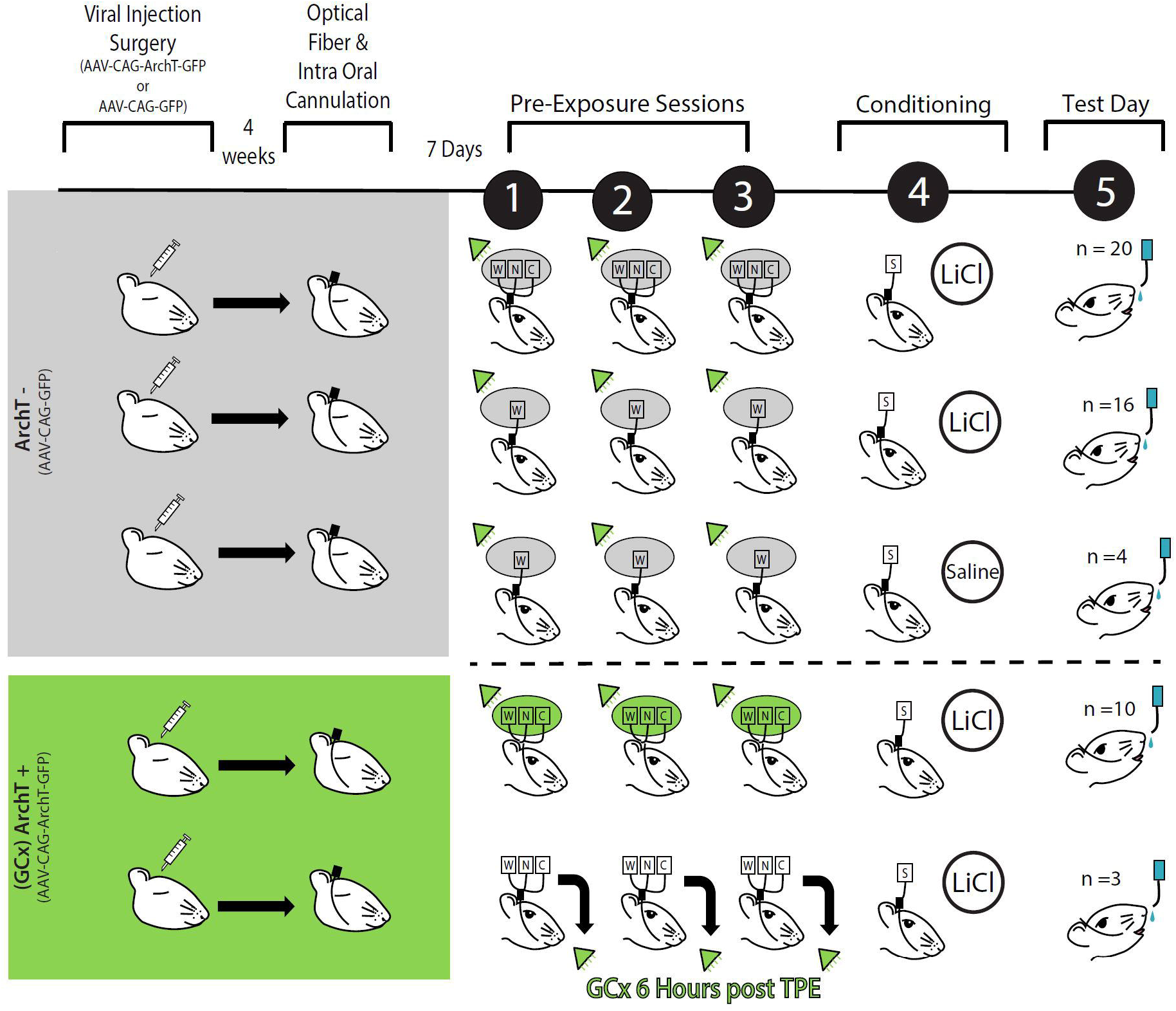
GCx during Taste Pre-Exposure paradigm. A complete timeline of the optogenetic (Experiment 2A) paradigm showing all groups. Rats first undergo viral injection surgery (either AAV-CAG-GFP or AAV-CAG-ArchT-GFP infused bilaterally into GC). Rats receiving AAAV-CAG-ArchT-GFP (ArchT+) are highlighted in green. Rats receiving only AAV-CAG-GFP (ArchT-) are highlighted in gray. To ensure high levels of viral infection, the optical fiber and intra oral cannulation surgery took place 4 weeks after viral injection surgery. Following 7 days of recovery after the optical fiber and intra oral cannulation surgery, all rats encountered 3 TPE (or WPE) sessions, with 532nm laser illumination of GC during each fluid exposure (indicated by green triangle). ArchT-groups (top 3 rows) involved 3 groups: from the top, TPE followed by sucrose + LiCl, WPE followed by sucrose + LiCl and WPE followed by sucrose + saline. ArchT+ groups (bottom 2 rows) involved 2 groups: TPE followed by sucrose + LiCl and TPE followed by sucrose + LiCl in which laser illumination was delayed by 6 hours (see text). On the day following conditioning, aversion strengths are tested via sequential presentation of sucrose and water for all 5 groups.

Rats in three of the 5 groups were injected with a “control virus” carrying only GFP (“ArchT-” groups–GC was illuminated by the laser, but this did not cause GC neurons to be inhibited): 1) an ArchT-group that received TPE followed by CTA training (n = 20); 2) an ArchT-group that received WPE followed by CTA training (n = 16); and 3) an ArchT-group that received WPE followed by sham conditioning (n = 4; Figure 6).

Running these control groups allowed us to replicate the basic, previously-reported (Flores et al. 2016) effect of TPE on learning in a new context: CTA in surgically-prepared, laser-illuminated ArchT-rats is enhanced (that is, consumption of newly-aversive sucrose in the testing session is reduced) by TPE (left WPE and TPE bars, Figure 7A).

**Figure 7.**
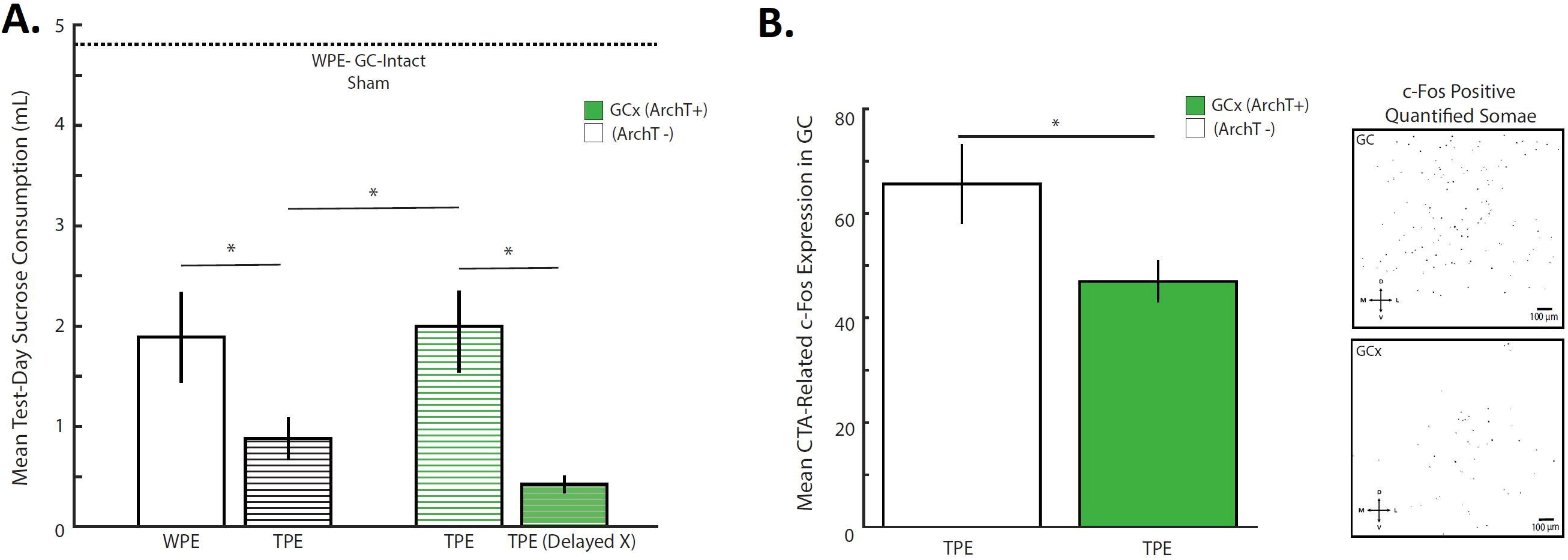
GCx during TPE inhibits CTA learning enhancement and impacts CTA-related c-Fos. **A.** In ArchT-rats (black and white bars, left side of graph), aversions to novel sucrose were stronger following TPE (black striped bar) than following WPE (open bar)—a replication of the original behavioral result (Flores et al. 2016). ArchT+ rats receiving GCx during TPE (thin green striped bar) showed significantly weaker CTAs compared to identically run ArchT-rats (compare the middle pair of TPE bars)—demonstration that GC activity during TPE is vital for the behavioral phenomenon. GCx induced 6 hours after each TPE session (thick green striped bar) did not reduce aversion strength. Finally, all conditioned groups showed learning when compared to ArchT-sham-conditioned rats (horizontal dashed line). The x-axis represents average raw sucrose consumption on test day (mL) across all groups. **B. *Left:*** CTA-related c-Fos expression in GC was significantly stronger in ArchT-rats (open bar), compared to ArchT+ rats (green bar)—GCx reduced CTA-induced c-Fos. ***Right:*** Representative images of c-Fos positive somae (masked in black) for GC-intact (top) and GCx (bottom) rats, quantified by the FIJI Analyzing Particles tool. Error bars represent SEM. (*) *p* < 0.05.

The two additional groups were infected with the AAV containing ArchT (GCx rats, Figure 6) such that laser illumination of GC inhibited neural firing. For one of these groups, GC neural inhibition (n = 10) was targeted to the period from 0.85 sec before until 2.5 secs following each intraoral taste infusion of each TPE session (i.e., the same parameters used with ArchT-rats).

These data allowed us to confirm our most central Experiment 2A hypothesis (Figure 7A), specifically that GCx during TPE effectively blocks the expected enhancement of CTA. That is, the strength of conditioning when GC neural activity had been disrupted during TPE (Figure 7A, 3^rd^ bar) was less than the strength of conditioning when GC was intact during TPE (Figure 7A, 2^nd^ bar), and was similar in strength to learning of WPE rats (Figure7A, 1^st^ bar).

While the absolute degree and duration of GCx was quite small (GC activity was suppressed for a total of 5.6 min in each of three 25-min sessions, and for only 3.35 sec at a time), we ran an additional control for the remote confound that GCx itself (as opposed to inhibition of GC during the presence of tastes) might reduce later learning. In this control condition, we induced GCx (n = 3; ArchT+) using precisely the same parameters as those described above (3.35 sec GCx every 15 sec, for a total of 5.6 min), but doing so hours after the end of each actual TPE session (see the Methods and Discussions sections for logic explaining why this specific protocol was used, rather than one in which GC was inhibited during TPE sessions but between taste deliveries).

This protocol utterly failed to diminish the TPE-related enhancement of learning (Figure 7A, rightmost bar), thus demonstrating that it is specifically GC activity during the TPE session that is important for the learning enhancement.

Statistical analysis of the data shown in Figure 7A confirmed each finding described above. A one-way ANOVA revealed that test day sucrose consumption was different across the 5 optogenetic groups (*F* (4, 31) = 13.867, *p* = 0.000). Fisher’s LSD post hoc analyses confirmed replication of the original enhancement of learning caused by TPE, in that learning was significantly enhanced (i.e., test day sucrose consumption was reduced) when CTA was preceded by TPE (0.879 ± 0.212 mLs) compared to WPE (1.890 ± 0.450 mLs; Fisher’s LSD *p* = 0.032) for ArchT-rats receiving laser illumination during pre-exposure trials.

More centrally regarding the current inquiry, this TPE-driven learning enhancement was blocked by GCx during TPE: learning was significantly stronger (i.e., test day sucrose consumption was lower) in ArchT-rats in which CTA had been preceded by TPE (0.879 ± 0.212 mLs) than in identically treated ArchT+ rats (2.056 ± 0.374 mLs, Fisher’s LSD *p* = 0.009). Inhibition of GC during TPE specifically impacted later learning, reducing it almost precisely to that observed with WPE (compare the 1^st^ and 3^rd^ bars of Figure 7A).

Lastly, our results clearly rule out the confounding possibility that inhibition of GC activity at a time in which tastes are not being presented impacts future processing: test day sucrose consumption was lower in ArchT+ rats for which GCx occurred 6 hours post TPE sessions (0.423 ± 0.084 mLs) than in rats for which GCx occurred during TPE (2.056 ± 0.374 mLs, Fisher’s LSD *p* = 0.015, Figure7A); it was if anything, even lower than consumption in unperturbed TPE rats (although it is important to note that this difference failed to reach significance, and therefore is not discussed further).

Rats in all 4 trained groups consumed far less than rats receiving TPE followed by a sham-training (horizonal line in Figure 7A, 4.808 ± 0.042 mLs; Fisher’s LSD *p* = < 0.05 for all comparisons), proof that our experimental paradigm was able to successfully implement aversions in AAV animals. Together these results strongly imply that GC processing of tastes during TPE is vital for the enhancement of learning normally observed after TPE.

Ancillary analysis failed to reveal anatomical specificity of the GCx effect. While fiber placements were too consistent to allow for an analysis of differences in the anterior-posterior plane, Pearson’s correlations (these data were normally distributed) showed no reliable dependency between average depth of fiber placement (see Methods) and raw test day sucrose consumption, within the TPE (*R* = 0.156, *p* = 0.648), WPE (*R* = 0.405, *p* = 0.191) or TPE-GCx (*R* = 0.393, *p* = 0.333) groups (Figure 5B; for other groups, N was too small to analyze).

In summary, the above results strongly support the view that GC activity during TPE changes how the brain handles the paired presentation of taste and LiCl in later CTA training. More specifically, they suggest that the elevations in CTA-related cortical c-Fos expression, observed in rats that had received TPE in the preceding days, may truly reflect an enhancement in the expression of learning caused by TPE.

#### Experiment 2B (impact of GCx on TPE-induced c-Fos)

Perhaps the best evidence for or against the theory supported above can be had from the testing of one additional, further prediction: if the above logic is correct, then GC perturbation during TPE, which inhibits the normal TPE-induced enhancement of learning, should also reduce the TPE-related enhancement of c-Fos expression.

We directly tested this prediction by comparing CTA-related c-Fos expression in rats that had undergone GCx during TPE to that observed in rats in which GC was unperturbed during TPE. ArchT+ (n = 9) and ArchT-(n = 8) rats underwent identical surgeries, identical TPE (with laser illumination synced with taste infusions), identical training protocols (pairings of taste and LiCl), and identical harvesting of brains 90 mins after the pairing (Figure 7B). The sole difference between groups was the nature of the virus injected prior to the onset of the protocol, and, thus, the impact of laser illumination (Figure 6).

Figure 7B presents the results of this experiment, which support our prediction. While overall c-Fos levels were lower in this experiment than in Experiment 1(a result that likely reflects the trauma of surgery, virus infusion, and ferrule implantation), GCx during TPE reliably reduced levels of c-Fos expressed in response to later CTA training, compared to rats for which TPE was offered with GC intact (*t* (15) = −2.238, *p* = 0.041). In fact, the reduction in c-Fos expression caused by GCx here is similar to the difference between TPE and WPE in Experiment 1.

Consistent with the findings in Experiment 1, anatomical analysis using two-way repeated measures ANOVA revealed absolutely no dependency of AP zone on this effect — GCx during TPE reduced training-induced c-Fos similarly for anterior, middle and posterior regions of GC. These results take our experiments full circle, demonstrating that GC plays a role in both incidental taste experience processing and taste association learning. Activity in GC during TPE processes the taste experience, allowing later enhancement of GC processing of CTA training, and thereby strengthening learning itself.

## DISCUSSION

In the current report, we have addressed the question of how non-reinforced incidental taste experience influences the neural processes underlying later learning. Such investigations are potentially of great importance, both because incidental taste experience is omnipresent in the natural world (and, thus, constantly exerting a noticeable impact on brain and behavior) and because the work reveals limits of the generalizability of data collected from laboratory animals lacking such experience.

Our data reveal that innocuous taste experience specifically affects processing of the association between taste and malaise in gustatory cortex (GC). TPE enhances conditioning-session induction of the immediate-early gene c-Fos (expression that both reflects activity levels post aversion learning and has been shown to be pivotal for the formation of CTA enduring memory; see Swank and Bernstein 1994; Navarro et al. 2000a; Spray et al. 2000; Koh and Bernstein 2005; Wilkins and Bernstein 2006; Yasoshima et al. 2006; St. Andre et al. 2007; Doron and Rosenblum 2010; Uematsu et al. 2015; Mayford and Reijmers 2016), while having no impact on the processing of a new taste (sucrose, see below) or nausea alone. We also found no significant differences in the expression of c-Fos across the anterior to posterior axis of the structure, suggesting that, while distinct sub-divisions of GC may contribute differently to CTA memory itself (Schier et al. 2016), TPE affects learning-related activity uniformly across the entirety of GC.

We went on to show that GC neural activity during TPE plays an indispensable role in enhancing CTA memory. GCx during TPE completely suppressed the TPE-induced enhancement of learning (whereas an identical GCx protocol delivered 6 hours after TPE sessions had no such impact), as well as suppressing TPE-induced enhancements of learning-related c-Fos. These results suggest that GC network activity elicited by innocuous taste experience may impact later learning by specifically priming the association between the novel taste and malaise.

### Learning-related activity in GC does not suggest TPE-induced changes in sucrose processing

In associative learning (of which CTA is a popular model), memory strength is influenced by the salience (or novelty) of both the CS and US—in the case of CTA, these are the taste (Lubow and Moore 1959; Ahlers and Best 1971; Logue 1979; Franchina and Slank 1988; Siddle et al. 1989; Rosenblum et al. 1993; De La Casa and Lubow 1995; Merhav and Rosenblum 2008; Clark and Bernstein 2009; Lin et al. 2012c) and malaise (Revusky 1968; Nachman and Ashe 1973; Flores et al. 2016; Levitan et al. 2016b)—as well as by the degree of their association (Garcia et al. 1966; Revusky 1968; Nachman 1970; Ahlers and Best 1971; Kalat 1971; Adaikkan and Rosenblum 2015). Expression of c-Fos in GC is similarly dependent on stimulus salience (Koh et al. 2003; Lin et al. 2012c). The impact of TPE on learning, and on c-Fos expression in GC, could therefore conceivably reflect an increase in salience of either stimulus: if TPE increased the novelty of novel sucrose, this change in sensory coding could drive the observed CTA changes, thereby implicating the cholinergic system, manipulations of which are known to be linked to taste novelty (Miranda et al. 2000; Miranda et al. 2003) and novelty-related CTA changes (Gutiérrez et al. 2003; Clark 2009; Neseliler et al. 2011), in TPE phenomenology.

Our analysis of c-Fos, consumption and licking reveal this explanation to be highly unlikely, however. It is well established that administration of novel tastes (including those that cause neophobic reactions) causes higher c-Fos induction than familiar tastes (Koh et al. 2003; Lin et al. 2012a); the fact that neither sucrose nor LiCl exposure-alone caused stronger c-Fos induction in TPE rats than in water-exposed rats suggests that TPE did not increase sucrose novelty or LiCl salience. In a separate brief access task experiment known to be particularly sensitive to novelty, TPE rats did not consume more sucrose than WPE rats—analysis of neither a 15-min exposure nor even of the initial 3 min of the 15-min session revealed any evidence of enhanced novelty/neophobia in consumption or licking of TPE rats. We conclude that TPE specifically affects cortical activity related to the central process involved in associative learning—i.e., the association between the stimuli—rather than the processing of taste or malaise alone.

Note that we are not saying that GC is not involved in the processing of sucrose (it almost certainly is); we are saying only that our c-Fos and behavioral data suggest that TPE’s impact is largely limited to enhancing the processing of the taste-malaise association—the learning effect is not secondary to a significant enhancement of taste novelty/neophobia. Our work does not therefore provide specific evidence suggesting involvement of the cholinergic system (known to be involved in taste novelty, see (Miranda et al. 2000; Miranda et al. 2003) in TPE.

### GC plays a specific role in mediating the impact of TPE on CTA learning

Of course, the above results do not prove a specific role for GC activity during TPE, either. The c-Fos data do not directly test whether TPE-induced changes in GC activity are truly important for CTA learning; other CTA-relevant brain regions that are connected to the GC, notably including the basolateral amygdala (Allen et al. 1991; Grossman et al. 2008; Matyas et al. 2014), could potentially mediate the effect on c-Fos expression during CTA. To more directly test the importance of GC activity during TPE, we made use of the optical silencer Archaerhodopsin-T (ArchT, see Yizhar et al. 2011; Maier et al. 2015; Li et al. 2016), and found that temporally controlled perturbation of GC during TPE reduces aversions to a strength similar to that observed in rats pre-exposed only to water (see Flores et al. 2016). Neither laser illumination alone nor GCx administered after TPE sessions hindered the TPE-related enhancement of CTA strength. These results strongly suggest that GC plays a role in the integration of TPE into future taste learning.

The fact that GC activity during TPE is important to CTA learning suggests a causal relationship between TPE activity and training-related c-Fos induction. We performed one further set of experiments to test this implication, showing that GC activity during TPE is important for not only TPE-induced enhancement of CTA, but also for the attendant enhancement of GC activation—GCx during TPE resulted in lower GC CTA-related c-Fos expression. Taken in context with the above, these results suggest that GC activity during “innocuous” taste experience promotes CTA learning by specifically priming the degree of association between taste and malaise. These novel findings are important since they implicate GC activity in the processing of incidental taste experience into future associative learning.

We cannot yet say what cortical process is being interrupted by our optogenetic inhibition. The fact that identical amounts of inhibition delivered a few hours after each TPE session had no deleterious impact on learning confirms that the effect likely reflects perturbation of the processing of the tastes themselves and not perturbation of a process lasting hours (Kandel et al. 2014; Levitan et al. 2016a); it remains possible, however, that the seconds to minutes after taste administration are important for a consolidation process that must occur following a taste experience. If so, perhaps inhibition induced immediately after TPE sessions (or a more molecular perturbation of plasticity such as protein synthesis inhibition, e.g. Ferreira et al. 2005; Garcia-DeLaTorre et al. 2009; Inberg et al. 2013; Levitan et al. 2016a) or even between taste deliveries would forestall the CTA enhancement observed here. It would be fascinating and valuable to test the time-course of this phenomenon across hours, as has been done in recent work on CTA consolidation (e.g., Levitan et al. 2016a).

These experiments were not done here, however, because their results would be difficult to interpret without a great deal of additional experimentation: if between-trial perturbation proved to have an impact on later learning, for instance, it would be necessary to ask whether that impact reflects a simple need for GC patency during TPE sessions, or whether taste processing itself, which is known to extend well beyond the period of taste delivery (Yamamoto et al. 1985; Katz et al. 2001; Katz et al. 2002), continues throughout a 30-sec interval. Similarly, an impact of inhibition during water pre-exposure could reflect functioning of a broader process, or could reflect the fact that water is itself a taste (Rosen et al. 2010), and thus that water pre-exposure is simply a weak, 1-taste TPE (Flores et al. 2016). Thus, we leave these necessarily large sets of experiments for later investigation.

Although also beyond the scope of our current work, we can begin to speculate regarding possible sources of the TPE effect. It is likely that innocuous taste exposure becomes associated with safe outcome (Lubow and Moore 1959; Ahlers and Best 1971; Lubow 1973; Lovibond et al. 1984; De La Casa and Lubow 1995; Lubow 2009; Monk et al. 2014). The multiple sessions of multi-taste pre-exposure that make up TPE would strengthen this association, and perhaps even leave rats with a general belief that “tastes are safe.” The subsequent pairing of a novel taste with an aversive outcome (during CTA) would saliently deviate from this expectation, a fact that would result in enhanced learning. This speculation is consistent with the fact that CTA learning enhancement following TPE is correlated with number of tastes to which the rat is safely pre-exposed (Flores et al. 2016).

### Possible cellular mechanisms

At a more mechanistic level, the fact that TPE influences learning that occurs days later suggests a metaplastic mechanism—activity-dependent priming of future cellular plasticity (Parsons 2017). One study examining this phenomenon showed that experience with one learning paradigm (olfactory learning) enhanced CA1 excitability in a manner that later correlated with enhanced learning of a different learning task (water maze; Zelcer et al. 2006). It is possible that TPE induces similar metaplastic changes—perhaps at synapses linking basolateral amygdala to gustatory cortex (BLA→GC), which are associated with CTA learning and have been shown to undergo metaplastic changes (Grossman et al. 2008; Rodriguez-Duran et al. 2011). Perhaps GC activity during TPE reduces the threshold for the induction of (BLA→GC) synaptic plasticity in future CTA conditioning sessions, thus enabling the enhancement of CTA learning (most likely via NMDA-dependent mechanisms, see Escobar and Bermudez-Rattoni 2000; Ferreira et al. 2005).

Particularly relevant guidance for a future investigation of molecular mechanism comes from work examining varieties of pre-experience and learning in several behavioral tasks including CTA (Ballarini et al. 2009; Moncada et al. 2011). It has been shown, for instance, that protocols that normally fail to produce lasting CTA memories may be bumped “above threshold” by previous (or, intriguingly, later) exposure to a different taste stimulus that can provide the necessary plasticity related proteins (Merhav and Rosenblum 2008; Ballarini et al. 2009); while this phenomenon, dubbed “behavioral tagging,” differs in several important ways from the phenomenon that we report here [in addition to using a sub-threshold learning protocol, in “behavioral tagging” the effective pre-exposure is a novel stimulus presented as little as 1 hr. prior to training, whereas the TPE effect is stronger following multiple presentations in the days prior to training; see (Flores et al. 2016 for more discussion)] it is intriguing to think that TPE stimuli could induce plasticity related genes (Inberg et al. 2013; Inberg et al. 2016) that are similar to those identified in this earlier work, could act in the same way as behavioral tagging.

Protein synthesis in GC itself has also been identified as vital for a related phenomenon, whereby latent inhibition learning (the ability of pre-exposure to the eventual CS to reduce CTA strength) is itself strengthened by “pre-pre-exposure” to a different, novel taste (Merhav and Rosenblum 2008). Again, this phenomenon differs from ours in many ways, but it would be unsurprising if in both paradigms protein synthesis driven by presentation of putative “innocuous” tastes was enhancing the next learning opportunity to arise. Future experiments will pursue the nature—and, just as importantly, the time course—of such protein synthesis.

Regardless of the ultimate underlying mechanism, our findings suggest a causal relationship between TPE activity in GC and the strength of later taste aversion learning-related activity. GC activity during “innocuous” taste experience promotes CTA learning by specifically priming the degree of association between a totally novel taste and malaise. These findings are important in that they grow our currently meager understanding of the neural substrates underlying the integration of taste experience with future associative taste aversion learning, a subject with great relevance to human research, and do so in a quite surprising way: they demonstrate that primary sensory cortex, far from being just involved in perception or even learning, is involved in integrating recent unreinforced experience with current stimulus associations. Our results highlight the need for future research into precisely how seemingly innocuous experience affects future learning at both the behavioral and neural levels.

## MATERIAL & METHODS

### Subjects

127 naïve adult (6-8 weeks, 225-250g at time of surgery) female Long Evans rats acquired from Charles River Laboratories (Wilmington, MA) served as subjects for all experiments. Females were chosen for their behavioral temperance, and because published evidence does not show major sex differences in CTA trainability (Randall-Thompson and Riley 2003; Rinker et al. 2008; Dalla and Shors 2009). Rats were housed individually in humidity-and temperature-controlled cages (Innovive), kept on a 12-hour light-dark cycle, and given *ad libitum* access to food and water (prior to experiments) which was replaced twice a week. At least 10 days post arrival to the facility, all animals were randomly assigned to experimental groups.

All procedures were conducted in accordance with the guidelines established by the Brandeis University Institutional Animal Care and Use Committee (IACUC).

### Stimuli

Taste solutions used for taste pre-exposure sessions (TPE) consisted of 0.01M sodium chloride (N) and 0.02M citric acid (C), as well as distilled water (W). Both concentrations are comfortably above detection thresholds for rats (Hiji 1967; Kolodiy et al. 1993; Scalera 2004; Li et al. 2012; Sadacca et al. 2012). These specific stimuli were used to ensure experience with both palatable and less palatable tastes—this concentration of N is modestly palatable, and this concentration of C is mildly aversive. A novel, innately palatable 0.2M sucrose (S) solution was always used as the conditioned stimulus in all experiments to facilitate detection of aversion learning. Note that while this means that we did not counterbalance completely, for instance making N the conditioned stimulus and S a TPE stimulus for some rats, in our previous work we demonstrated some basic generalizability, showing that N and C were similarly effective as single TPE stimuli (Flores et al. 2016).

### Experimental Apparatus

Experiments were conducted in the morning, following a 21-hr water deprivation period. All sessions occurred in a Plexiglass experimental chamber (8.5 × 9.5 × 11.5 in) that was distinct from the rats’ home cages. The experimental chamber and bottles were rinsed and sterilized before and after each use.

#### Experiment 1

##### Surgery

Rats were anesthetized using a ketamine/xylazine mixture (1ml ketamine, 0.05 ml xylazine/kg body weight) delivered *via* intraperitoneal injection. The head was shaved and positioned into a stereotaxic device, after which the scalp was exposed, leveled, and cleaned. Four self-tapping support screws were implanted into the skull.

Rats were then removed from the stereotax and laid prone for bilateral implantation of an intra oral cannula (IOC)—flexible hollow plastic tubes inserted parallel to the masseter muscle into the mouth posterolateral to the first maxillary molar (Phillips and Norgren 1970). A stable, rigid dental acrylic head cap was formed around the IOCs and skull screws.

Following surgery, rats were given analgesic (meloxicam 0.04 mg/kg), saline, and antibiotic (Pro-Pen-G 150,000 U/kg) injections. Additional antibiotic and analgesic injections were delivered 24 and 48 hours later. The weight of each animal was recorded each day; any rat displaying lethargy, lack of grooming or weight loss greater than 15% of pre-surgery weight were removed from the study. All rats were given 7 days of recovery post-surgery before any experimentation.

##### Adaptation sessions

Following recovery, rats were given 2 days of access to distilled water through a bottle in the experimental chamber for approximately 30 min. This ensured familiarization with the testing environment.

##### TPE/WPE sessions

TPE and water pre-exposure (WPE) sessions, which followed adaptation, were identical to those used in our previous experiments (Flores et al. 2016). Each rat (n = 66) received 1 such session per day—100 aliquots of either tastes (pseudo-random ordering) or water alone, delivered to the oral cavity *via* IOC (brief opening of a solenoid valve caused 50μl of fluid to be delivered) at 15-second inter-trial intervals, for a total of 5 mL of fluid. This protocol continued for 3 consecutive days (Figure 1). The IOC was used for delivery of TPE because it ensured experimenter control, such that TPE rats consumed equal volumes of unpalatable C and palatable N and WPE rats received the same volume of W.

##### Conditioning Sessions

A single conditioning session took place the day after completion of TPE/WPE, in the same experimental chamber. Sucrose CS was delivered *via* bottle (5mL available for 5 min) and then IOC (60 deliveries for a total of 3mL; Figure 1). This procedure allowed us both to: 1) take advantage of the literature indicating that c-Fos expression is stronger with bottle than IOC conditioning (Wilkins and Bernstein 2006) and 2) ensure substantial and reliable sucrose consumption. Immediately after sucrose consumption animals in the aversion conditioning groups (TPE, n= 15 and WPE, n = 15; Figure 1) received subcutaneous injections of lithium chloride (LiCl, 0.3 M, 0.5% of current bodyweight) to induce the malaise US. Use of this concentration of LiCl, which is lower than that typically employed to induce CTA, ensured that CTA learning would be sub-maximal, thereby allowing us to observe enhancements of learning (Nachman and Ashe 1973; Stone et al. 2005; Levitan et al. 2016b).

In addition to the above-mentioned WPE controls, additional control rats were given TPE followed by either: 1) S paired with subcutaneous injections of harmless saline—essentially S alone (n = 9); or 2) administration of LiCl alone, without pairing of S (n = 8; Figure 1). Finally, two further groups of controls received WPE followed by S + saline (n = 10) or LiCl-alone conditioning sessions (n = 9; Figure 1). LiCl/saline Injections were administered on the experimenter’s lap (a location distinct from both testing chamber and home cage), to ensure that malaise would not be associated with a context that could confound the results (or cause a great deal of potentially novelty-related c-Fos). Rats were then briefly (see below) returned to their home cages with access to *ad lib* food and water.

##### Brief Access Task

An additional set of animals (n = 8) took part in a brief access task (BAT, aka the “Davis rig,” Med Associates Inc.) to provide a separate, consumption test of the possibility that TPE changes sucrose processing itself.

The procedure was similar to previously used protocols to investigate neophobia (Lin et al. 2012c; Monk et al. 2014). Following IOC surgeries, recovery and adaption (see above), rats underwent 2 days of water habituation in the BAT rig, during which they learned to drink from the single lick spout in the chamber. Water deprivation for the 21 hours before each session ensured motivated drinking. At the start of each BAT trial, a mechanical shutter was raised to allow access to the lick spout for 15 mins (15mL of W), after which the shutter descended. The experimental chamber and bottles were rinsed and sterilized before use for each animal and session.

Following the two days of water habituation, animals underwent TPE (n = 4) or WPE (n = 4) sessions as described above. 24 hours after the last of these 3 pre-exposure sessions, they were returned to the BAT Rig, and this time given access to novel sucrose (15mLs available for 15 mins). Consumption, lick rate and interlick intervals were recorded for each rat. Rats were then returned to their home cages with access to *ad lib* food and water.

None of the rats in this experimental group (n = 8) were given CTAs toward sucrose or perfused for immunohistochemistry analyses.

##### c-Fos immunohistochemistry

To capture peak c-Fos expression levels (Chaudhuri et al. 2000), rats were deeply anesthetized with an overdose of the same ketamine/xylazine mix used for surgery ninety mins after the conditioning session and perfused transcardially with isotonic phosphate buffered solution (1X PBS) followed by 100ml of ice-cold 4% paraformaldehyde. Brains were then extracted and post fixed in 4% paraformaldehyde for three days, after which coronal brain slices (60 μm) containing the region of interest were sectioned on a vibratome. Sections were chosen based on anatomical landmarks for Gustatory cortex (GC (+2.5 mm for anterior GC, +1.4 mm for middle GC, +0.36 mm for posterior GC relative to bregma; see Paxinos and Watson 2007)) and on published reports demonstrating the regions importance to CTA or taste processing (Katz et al. 2001; Fontanini and Katz 2006; Piette et al. 2012; Sadacca et al. 2012; Maier and Katz 2013; Schier et al. 2014; Schier et al. 2016), the driving of taste-responsive behavior (Li et al. 2016; Sadacca et al. 2016), and the coding of taste learning (Stone et al. 2005; Fontanini and Katz 2006; Grossman et al. 2008; Moran and Katz 2014; Levitan et al. 2016a).

The c-Fos antibody protocol used was adapted from the manufactures recommendation (Santa Cruz Biotech: https://www.scbt.com/scbt/resources/protocols). Slices were rinsed with 1X PBS and incubated in a blocking solution (1XPBS/.3%TritionX-100/3% Bovine serum albumin) for 12 hours at 4^<u>°</u>^C. Blocking solution was removed and replaced with the primary antibody solution which consists of 1:100 c-Fos polyclonal rabbit IgG (SC-52G; Santa Cruz Biotechnology) for 12 hours at 4^<u>°</u>^C. After incubation, slices were rinsed using a 1XPBS/.3% Triton X-100 solution followed by the secondary antibody incubation of 1:500 c-Fos Alexa Flour 546 Goat-Anti-Rabbit IgG (H+L) (Life Technologies) and 5% natural goat serum for 12 hours at 4^<u>°</u>^C. Sections were then rinsed 5-6 times over 90 mins (1XPBS/.3% Triton X-100) and mounted on charged glass slides and cover slipped with antifade mounting medium with DAPI (Vectashield) to verify that c-Fos expression was specific to the nucleus of GC cells.

To monitor the expression of c-Fos, bilateral GC sections (limited to ventral gustatory cortex (Figure 2A & 4A) were viewed by confocal fluorescence microscopy with a Leica Sp5 Spectral confocal microscope/Resonant Scanner with 405 lasers equipped with *x*/*y*/*z* movement stage. Imaging and quantification was performed blind—the experimenter was unaware of the experimental group from which tissue was collected at the time of analysis.

#### Experiment 2

##### Virus Injection Surgery

Rats were anesthetized and prepped as for Experiment 1. With the skull exposed, cleaned, and leveled, bilateral craniotomies were then made at stereotactic coordinates above the part of GC (AP = 1.4 mm, ML = 5 mm from bregma; see Paxinos and Watson 2007) previously shown to contain neurons that respond distinctly to tastes in awake rats (e.g., Katz et al. 2001; Katz et al. 2002; Maier and Katz 2013; Sadacca et al. 2016).

We infused either adeno-associated virus (AAV serotype 9, n = 23) coding for ArchT and green fluorescent protein (AAV9-CAG-ArchT-GFP, 2.5×10^11^ particles/mL) or control virus coding for green fluorescent protein alone (AAV9-CAG-GFP, n = 13, 2.5×10^11^ particles/mL, http://www.med.unc.edu/genetherapy/vectorcore) into GC. Thus, we had “ArchT+” and “ArchT-” rats. ArchT has been shown to have better light sensitivity (1–10 mW/mm^2^) than other optogenetic AAV constructs, and thus to be particularly useful for investigations of the neural mechanisms of behavior (Zhang et al. 2010; Yizhar et al. 2011). This AAV serotype has been shown to effectively spread and infect all cell types (Aschauer et al. 2013), and to be effective within GC (Maier et al. 2015; Li et al. 2016).

Viral particles were suspended in a phosphate buffered solution containing Oregon Green 488 (Invitrogen). To infect GC, micropipettes (tip diameter 10-20um) carrying this solution were lowered to a sequence of three depths (4.9, 4.7, and 4.5 mm from dura), at each of which virus was delivered in discrete pulses (50 nl/pulse, 7 seconds between each pulse) controlled by an automatic Nanoject III Microinjector (Drummond Scientific). Following each unilateral set of injections, micropipettes remained in place for 5 min, and were then smoothly removed over the course of one minute so that fluid would not spread back up the cannula track.

A total volume of 1.25 μl of virus was delivered in 25 pulses per each injection depth. Following bilateral injections wounds were sutured and rats entered post-operative care. To ensure high expression, all viral injections detailed here were made 4 weeks prior to any further procedures. During these 4 weeks rats remained in their home cages, with no experimental manipulation.

##### Optical Fiber & Intra-Oral Cannulation surgery

Four weeks after viral injection surgeries, rats were again anesthetized and prepped for surgery. Following the insertion of four self-tapping support screws, the bilateral GC craniotomies created during the previous surgery were re-opened. Custom-built optical fiber assemblies (multimode fiber, 0.22 numerical aperture, 200 μm core, inserted through a 2.5mm multimode stainless-steel ferrule; THORLABS) were lowered to 4.7 mm ventral to the reflected dura mater (targeting the center of virus expression).

Placement of the fibers was stabilized with dental acrylic. Once the dental acrylic was dry, rats were removed from the stereotax and laid prone, and IOCs were implanted bilaterally (see above). The rigid dental acrylic head cap included the optical fibers, IOCs and skull screws. Rats were given post-operative care as detailed above, as well as 7 days or recovery prior to the start of experimentation.

##### Adaptation sessions

Following recovery, rats were given 2 days of adaptation, as *per* Experiment 1.

##### TPE/WPE sessions

Following adaptation sessions, rats were given either TPE or WPE sessions (1/day) for 3 days, as in Experiment 1. For Experiment 2, however, fluid deliveries were accompanied by optical illumination using 532nm laser light (Shanghai Dream Lasers), coupled to optical fibers (multimode fiber, 200 μm diameter, 0.22 NA) using customized FC/PC patch cables (THORLABS). For every taste delivery, the laser was turned on 850ms before a solenoid valve opened to release a taste onto the tongue and turned off 2500ms later (see Figure 5C; Mukherjee et al. 2017). Precisely this same protocol was run for 4 groups of both ArchT+ and ArchT**-** rats (see below), ensuring that: 1) GC activity was perturbed during the period in which we know taste processing to occur post-delivery (Katz et al. 2001; Katz et al. 2002); 2) GC was intact during the 15 second interval between all taste deliveries, as well as before/after each session; and 3) even control rats received GC virus infection and taste-coincident laser illumination.

The power of illumination was adjusted, before implantation, to be 40mW at the tip of the fiber using an Optical Power and Energy Meter console (THORLABS); this intensity has been calculated to inactivate neurons *in vivo* within an approximate 1mm sphere around the tip of the optic fiber (Han et al. 2011; Yizhar et al. 2011)—a sphere that encompasses about 33% of GC in the caudal-rostral axis (Kosar et al. 1986; Maier et al. 2015; Li et al. 2016). The same parameters have previously been shown to reduce the activity of ArchT+ single cortical neurons with minimal latency and damage (Maier et al. 2015; Li et al. 2016); pilot experiments (Figure 5C) revealed that the same is true in our rats.

In all, 5 groups were run (n = 53). Three of these were ArchT-groups—two TPE, and one WPE (allowing us to replicate the TPE effect in infected, ArchT-rats, and to evaluate consumption levels in sham-trained TPE rats). The remaining two were ArchT+ groups: 1) a group (the most important group) that allowed us to test the impact of GC inactivation during TPE; and 2) a group that received unperturbed TPE, and then, six hours after the taste session, received “delayed” but otherwise identical protocol of GC inhibition (allowing us to test the impact of GC inactivation lacking taste stimuli).

##### Conditioning Session

Conditioning sessions for Experiment 2A involved presentation of sucrose CS *via* IOC only (100 deliveries, 5mL total); this control ensure substantial and reliable sucrose consumption, such that otherwise-possible between-rat differences in amounts taste consumption could not confound the results (Figure 6). We have previously shown that both the Experiment 1 and 2 conditioning methods (i.e., Bottle + IOC; IOC-only) do cause taste aversions, although these aversions are somewhat stronger (as expected) with bottle delivery (Flores et al. 2016). Sucrose consumption was immediately followed by subcutaneous injections of lithium chloride (LiCl, 0.3 M, 0.5% of current bodyweight) to induce the malaise unconditioned stimulus. One group received sham training (sucrose + saline), and all others received CTA-inducing sucrose + LiCl.

In Experiment 2B, which explicitly examined the impact of cortical inactivation TPE on learning-related c-Fos expression, conditioning sessions were identical to those described for Experiment 1. No laser illumination was delivered during any conditioning session in Experiment 2A or B—that is, these experiments examined the impact of inhibiting GC activity during TPE sessions on aspects of conditioning delivered 1 full day after the end of the neural inhibition itself.

##### Testing Session (Experiment 2A)

Testing sessions were, like conditioning sessions, given without optical illumination. Rats were first presented with a bottle containing 5 mL sucrose (S) for 5 minutes; following a 5-minute pause in which no bottle was available, rats were then presented with a different bottle containing 5 mL water (W), again for 5 minutes.

Following test day, rats were deeply anesthetized with an overdose of the Ketamine/Xylazine mix and perfused as above. Brain tissue was harvested for localization of optical fiber placement.

##### c-Fos immunohistochemistry (Experiment 2B)

Rats used for Experiment 2B did not receive testing sessions. Instead, they were perfused for c-Fos analysis 90 minutes after the US administration in the conditioning session. Collection and analysis of immunohistochemistry proceeded as for Experiment 1.

##### GFP expression

To monitor expression of the AAV infection *via* visualization of GFP, slices were stained using previously developed protocol (Maier et al. 2015; Li et al. 2016). All slices were rinsed 3 times with 1XPBS over 15 mins. Slices where then permeabilized in a 0.3% Triton X-100/1% normal Donkey serum/1XPBS blocking solution for 2 hours at room temperature. Blocking solution was removed and replaced with primary antibody solution which consists of 1:500 anti-green fluorescent protein – rabbit IgG fraction (Life Technologies) for 12 hours at 4^<u>°</u>^C. After incubation, slices were rinsed 3 times over 15 mins using a 1XPBS followed by the secondary antibody incubation of 1:200 Alexa Flour 488 donkey anti-rabbit IgG (H+L) (Life Technologies) for 12 hours at 4^<u>°</u>^C. Sections were then rinsed 3 times over 15 mins using 1XPBS. Slices were then mounted on charged glass slides and cover slipped with Fluoromount Aqueous Mounting Medium.

Slices were imaged with a Keyence fluorescent microscope to confirm successful virus infection and optical fiber location after each animal. Although spread of virus does not itself define the area of perturbation, fluorescence confirms that the region of interest was infected and more importantly, that the optical fiber tip was localized within that infected region. Rats (n = 5) that did not have confirmed bi-lateral virus infection and/or incorrect position of the optical fiber were excluded from the study.

#### Quantification and Statistical analysis

All results were analyzed using SPSS and MATLAB. Significance across all experiments was defined as *p* < 0.05.

#### Experiment 1

##### c-Fos quantification and analysis

Measures of c-Fos were collected for 6 independent groups of animals (TPE: Sucrose + LiCl, Sucrose + saline, LiCl alone, WPE: Sucrose + LiCl, Sucrose + saline, LiCl alone). To minimize systematic bias, c-Fos counts were performed blind and semi-automatically, using FiJi (University of Wisconsin-Madison) software (Schindelin et al. 2012). Using the Analyze Particles function (after first rejecting particles outside the size range 10-infinity um^2 and circularity of 0_elongated polygon_ - 1.00_perfect circle_ as non-neural), a manual threshold (0-0.50%) was applied across all samples to differentiate between background and somae. Un-normalized soma counts were averaged across hemispheres and anterior, middle and posterior regions of GC resulting in one soma count per slice/rat. The mean soma counts for each of the 6 groups were used in all statistical analyses.

For Experiment 1, we tested the hypothesis that TPE specifically enhances learning-related c-Fos expression in GC. Analysis began with a one-way ANOVA but centered on *planned comparisons* between subsets of conditions. This analysis was deemed more appropriate than a two-way ANOVA, because the planned comparisons cut across traditional main and interaction affects: our hypothesis specifically turned on an evaluation of whether CTA-related c-Fos activity when preceded by TPE was greater than that when: 1) CTA was preceded by WPE; 2) presentation of S alone was preceded by TPE; or 3) presentation of LiCl alone was preceded by TPE. Data passed assumptions of homogeneity of variances (Levene’s test, *p* = 0.118) and is presented as mean ± standard error.

The enhancement of CTA-related c-Fos expression caused by TPE was subject to further analysis, including a two-way repeated measures ANOVA that allowed us to evaluate any potential anatomical specificity of the phenomenon within sub-regions of GC. Again, data passed assumptions of homogeneity of variances (Levene’s test, *p* > 0.05 for all groups) and is presented as mean ± standard error.

##### Consumption analysis

To test the potential impact of TPE on sucrose consumption during conditioning sessions, we performed several analyses (using independent samples t-tests to test for significance of differences) on BAT data—comparing raw consumption of sucrose between TPE and WPE groups, as well as lick rate and initial (1^st^ 3 minutes) lick rate.

#### Experiment 2

##### Analysis of CTA learning

Sucrose and water consumption (mL) on testing day was used to evaluate learning (less sucrose consumption = stronger learning). Consumption for different groups was compared using a one-way ANOVA followed by Fisher’s LSD *post hoc* analyses. Data passed assumptions of homogeneity of variances (Levene’s test, *p* = 0.123) and is presented as mean ± standard error.

##### Localization of Optical Fiber

We also performed a Pearson’s bivariate correlation to determine any relationship between placement of optical fibers (i.e., depth) and test day sucrose consumption: each fiber was given a score of 1-4 (ranging from 1: granular, 2: dysgranular, 3: agranular dorsal and 4: agranular ventral as defined by Paxinos and Watson 2007) for each hemisphere; these scores were then averaged across each animal for a single Fiber Depth Index where 1 was most dorsal insular cortex and 4 was most ventral insular cortex.

##### cFos Quantification and Analysis

Slices in Experiment 2B were analyzed for cFos expression as in Experiment 1 (see above).

## ACKNOWLEDGEMENTS

This research was supported by the National Institutes on Deafness and other Communication Disorders (R01DC006666 to D.B.K and F31DC015931 to V.L.F). The authors thank Jian-You Lin, PhD for consulting with us on the use of the BAT, Madeline Lefkowitz for her assistance in animal care and Joseph Wachutka for his help in designing and constructing an opto-trode system for simultaneous electrophysiology and optogenetics in GC.

